# Lightweight Reinforcement Algorithms for autonomous, scalable intra-cortical Brain Machine Interfaces

**DOI:** 10.1101/2020.12.08.416131

**Authors:** Shoeb Shaikh, Rosa So, Tafadzwa Sibindi, Camilo Libedinsky, Arindam Basu

## Abstract

Intra-cortical Brain Machine Interfaces (iBMIs) with wireless capability could scale the number of recording channels by integrating an intention decoder to reduce data rates. However, the need for frequent retraining due to neural signal non-stationarity is a big impediment. This paper presents an alternate paradigm of online reinforcement learning (RL) with a binary evaluative feedback in iBMIs to tackle this issue. This paradigm eliminates time-consuming calibration procedures. Instead, it relies on updating the model on a sequential sample-by-sample basis based on an instantaneous evaluative binary feedback signal. However, batch updates of weight in popular deep networks is very resource consuming and incompatible with constraints of an implant. In this work, using offline open-loop analysis on pre-recorded data, we show application of a simple RL algorithm - Banditron -in discrete-state iBMIs and compare it against previously reported state of the art RL algorithms – Hebbian RL, Attention gated RL, deep Q-learning. Owing to its simplistic single-layer architecture, Banditron is found to yield at least two orders of magnitude of reduction in power dissipation compared to state of the art RL algorithms. At the same time, post-hoc analysis performed on four pre-recorded experimental datasets procured from the motor cortex of two non-human primates performing joystick-based movement-related tasks indicate Banditron performing significantly better than state of the art RL algorithms by at least 5%, 10%, 7% and 7% in experiments 1, 2, 3 and 4 respectively. Furthermore, we propose a non-linear variant of Banditron, Banditron-RP, which gives an average improvement of 6%, 2% in decoding accuracy in experiments 2,4 respectively with only a moderate increase in power consumption.

## I. Introduction

It is estimated that around 1 in 50 people worldwide are afflicted with paralysis [1]. Intra-cortical Brain-machine interfaces (iBMIs) are essential to restore quality of life for patients suffering from debilitating paralytic conditions such as tetraplegia, quadriplegia, stroke induced paralysis among others. Neural spikes recorded from the surface of brain areas associated with movement (e.g. primary motor cortex - M1) serve as an input to iBMI systems. The effector outputs vary depending on the application. For e.g., in order to enable communication, real-time demonstrations such as [2]–[4] employing a cursor to type out messages have been exhibited. Likewise for locomotion, real-time demonstrations in the form of wheelchair [5], full-body exoskeleton [6] have been reported.

Notwithstanding the impressive advances made in the field of iBMIs, practical challenges persist in clinical translation. Firstly, most iBMI systems use wired connection to bulky computers reducing patient mobility and increasing risk of infection [8]. Wireless iBMIs suffer from scalability issues due to exploding data rates [9]. A potential solution is including an intention decoder in the implant [10]; however, the need for frequent retraining makes it impractical.

Secondly, the current state of the art iBMI systems involve time-consuming daily calibration procedures amounting to several minutes, at the beginning of a given day/session [11]. This greatly affects the ability of iBMI users, suffering from debilitating impairments, to remain alert during these prolonged calibration routines [11]. Imagine having to calibrate your mobile touch screen every time before using your handset.

Lastly, these systems employ supervised learning algorithms to learn the mapping from input neural signal to output effector movement [12]. The supervised learning paradigm requires a ground truth target label to be available at every instance of time to train an appropriate model. In case of patients who cannot move their limbs to control effector kinematics, getting this ground truth target label requires some workaround. It involves carefully designed calibration trials that are designed with the assumption that the ground truth target label is always pointing towards the direction of the eventual goal in the designed trial [13], [14]. These requirements often necessitate the supervision of a neural engineer to ensure smooth operation of the iBMI system.

To tackle the above problems, we propose applying online reinforcement learning based methods that, (a) learn the neural input to effector movement mapping on incoming samples on a sample-by-sample basis, thereby getting rid of explicit calibration procedures and, (b) do not require explicit measurement of effector kinematics and learn with a simple scalar reward that is obtained while interacting with the external environment.

Briefly put, a reinforcement learning (RL) system involves an RL-agent outputting an *action* with inputs as - *states* and *rewards* at a given time-step - *t* = *i* (see Fig. 1(c)). *States* in case of iBMIs correspond to the input neural feature vector sensed from micro-electrode arrays at *t* = *i*, whereas *rewards* are scalar values obtained as a result of *action* taken at the previous time-step - *t* = *i* − 1. Maximizing the score of total reward is the objective of learning process in RL-based systems.

**Fig. 1:**
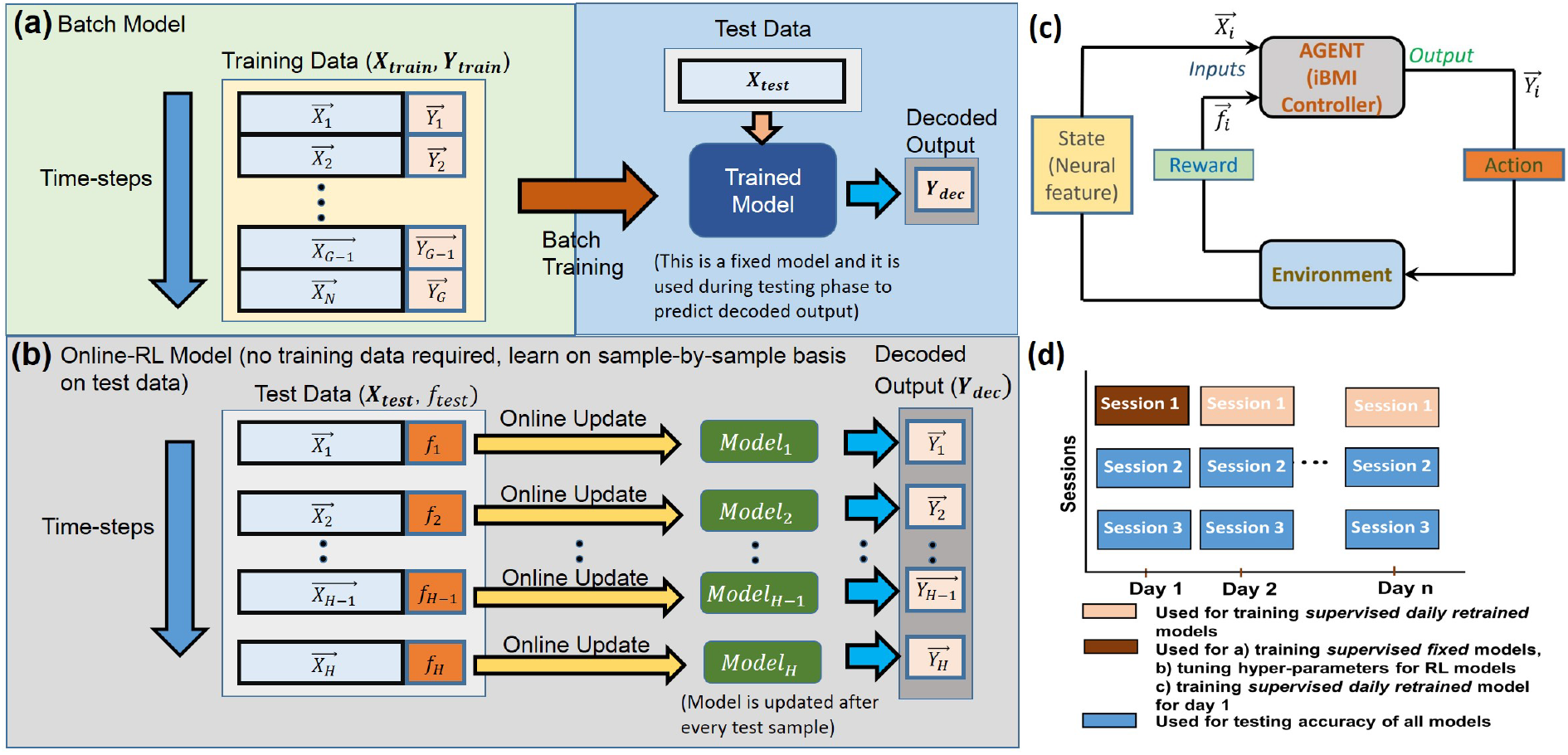
Shows (a) Batch and (b) Online Reinforcement Learning based learning schemes. Batch-trained models learn a model on an explicit set of training data. Typically, supervised learning schemes employed in iBMIs employ batch-based learning schemes. Online reinforcement learning-based models [7] do not employ training data for learning. Instead, they learn during the test phase by virtue of a scalar feedback 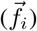 at every time-step, *t* = *i*. (c) shows block level representation of Reinforcement Learning in context of iBMIs. The iBMI controller emits a discrete output action 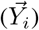 based on inputs - Neural states 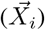 and rewards 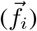 at every time-step, *t* = *i*. (d) shows the train-test data split. The first session’s data of every recorded day was used for training *retrained* supervised models across all pre-recorded datasets. First session’s data of the first day of recording was used to build *fixed* supervised models as well as tune hyper-parameters associated with Reinforcement Learning (RL) models. Sessions 2 and 3 of each day are used as test data across all techniques.

Promising RL based iBMI implementations have been reported in the past. This work aims to build up and extend on the reported techniques. Popular RL algorithm of the AlphaGo fame [15], Q-learning was reported in [16] wherein rats controlled a prosthetic arm to choose between two targets. The limitation of this approach is that the large neural state-action pair gives rise to the curse of dimensionality problem leading to generalization difficulty. To overcome this problem, [17] proposed applying Attention-gated reinforcement learning (AGREL) and its variants [18], [19]. [17] reports improvement over Q-learning involving one NHP performing a center-out cursor task. An important point to bear in mind is that the reported approaches trained the RL-models in batches implying that these methods would still require an explicit block of training similar to the state of the art supervised methods. Alternatively, [7] has presented an iBMI based on Hebbian Reinforcement learning (HRL) algorithm that is trained in an online fashion at each time-step. This allows us to get rid of an explicit calibration block and results have been reported in two NHPs performing a two-target discrete task [7]. Furthermore, [7] has introduced a simplified version of a binary scalar reward to train the RL-model and preliminary results have reported biological sources to contain this reward information [20], [21]. However, all the reported approaches use multilayer neural networks that are prone to the local optimum problem [19].

Besides generalization issues, multi-layer neural networks based approaches are found to be too resource intensive to be deployed in an implantable fashion [8], [22]. To overcome the issues of generalization and implantability, we present application of Banditron [23] - a lightweight single-layer online prediction algorithm in an iBMI setting. Furthermore, we propose and present a non-linear variant of Banditron, Banditron-RP (Random Projection, see Section III-A2) alongside its low power custom implementation.

Thus, in summary, we are presenting an alternate online BMI paradigm along the lines of [7], wherein we do not explicitly collect training, validation data (see Fig. 1(a)). Rather, we propose to train our model sequentially with incoming data on a sample by sample basis as seen in Fig. 1(b). This helps in getting rid of several minutes worth of fatigue-inducing calibration routines. In the experiments reported in this paper, the time required to conduct calibration routines ranges from approximately 5 to 14 minutes. The quantum of calibration time amounts to approximately one-third of the total experiment time in our reported experiments.

In keeping with this line of thinking, each incoming feature sample 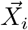 at time-step *t* = *i* is used to update the associated weights only once, based on a scalar feedback 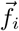. This methodology is consistent across all the reported RL algorithms.

The main contributions of this paper include

- Comparing the computational complexity of Banditron and Banditron-RP against state of the art RL algorithms - AGREL, HRL and Q-learning
- Comparing offline discrete-state decoding performance of Banditron and Banditron-RP against state of the art RL algorithms across four datasets spanning multiple days of recording
- Investigating the impact of erroneous and sparse nature of reward signal on the above reported RL algorithms
- Reporting RL algorithms decoding performance against popular supervised classification methods - Linear Discriminant Analysis (LDA) and Support Vector Machines (SVM). Do note that this is not a one-to-one comparison as RL and supervised methods employ altogether different paradigms of learning (see Fig. 1(a),(b)).

## II. Materials and Methods

### A. Signal Acquisition and Processing

We have used two adult male macaques (*Macaca fascicularis*) to conduct our experiments. We will refer to them as non-human primate (NHP) A and NHP B respectively. A titanium head post (Crist Instruments, MD, USA) was affixed prior to implantation of microelectrode arrays in both NHPs. In NHP A, 4 microelectrode arrays containing 16 electrodes each [5], and in NHP B 1 microelectrode array containing 100 electrodes were implanted in the hand/arm region of the left primary motor cortex respectively [8], [24]. Threshold crossings from the sensed neural signals were used as inputs [5], [25].

### B. Behavioural Tasks

We have collected a total of 4 datasets, two from NHP A (experiments 1 and 3) and two from NHP B (experiments 2 and 4) respectively. Pictorial representation of the tasks can be seen in Fig. 2. Brief description is provided below

**Fig. 2:**
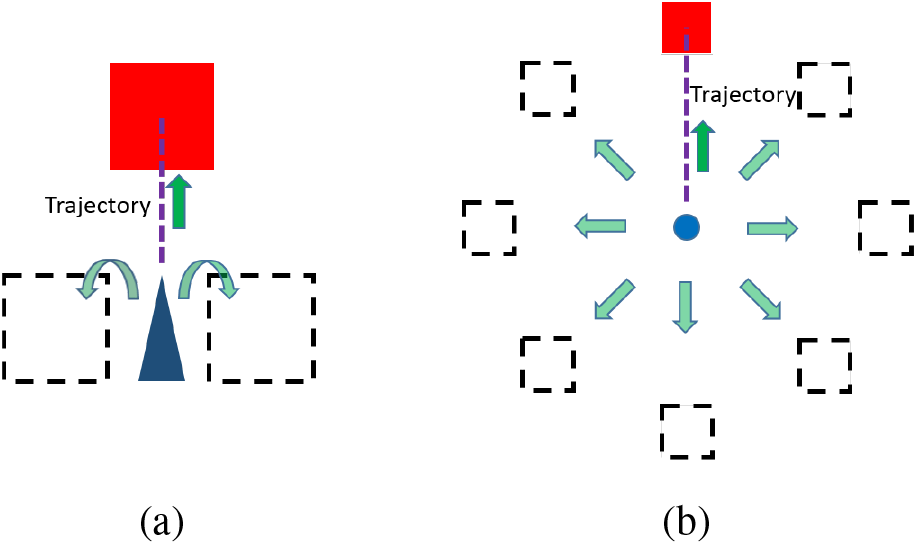
(a) This cartoon represents the general nature of the behavioural task performed in experiments 1, 2 and 3. In experiments 1 and 2, the nonhuman primate (NHP) is trained to maneuver a joystick-controlled wheelchair whose position updates every 100 ms. The wheelchair starts from a central position in every trial to reach the target over several time-steps, appearing in one of the three square-shaped locations [5]. However, in experiment 3, closed loop brain control was employed and decoded commands from a discrete classifier were used for controlling wheelchair motion. Wheelchair in both scenarios of brain and joystick control was driven at a frequency of 10 Hz. (b) represents a center-out task corresponding to experiment 4, wherein the NHP was trained to move a joystick-controlled cursor starting from a central position on a computer screen to one of the eight different square-shaped target locations. Cursor position was updated every 100 ms and the NHP reached the target over several time-steps.

#### 1) Experiment 1

This experiment involved a robotic wheelchair bound NHP A controlling its motion through a three-directional spring-loaded joystick [5]. The experiment comprised of four tasks - a) turning 90° right, b) moving forward by 2 m, c) turning 90° left and d) staying still for 5 seconds (*stop* task). A trainer holding a piece of reward served as a visual target and a trial was considered successful if NHP A managed to reach the trainer within an elapsed time of 15 seconds while the wheelchair was driven at a frequency of 10 Hz.

#### 2) Experiment 2

In this experiment, NHP B was seated in a primate chair facing a computer screen and was trained to manipulate a three-directional joystick similar to experiment 1. This experiment consisted of three tasks, with a target being presented in a pseudo-random manner in one of the following locations – Top, right and left relative to the centre of the screen. The target was in the form of a red square of side 2 cm on a black background. Each trial began with a wheelchair avatar being displayed at the centre of the screen along with the target. A trial was considered successful if NHP B managed to reach and stay in the target area for under a total elapsed time of 13 seconds. A juice reward was dispensed for every successful trial and the wheelchair avatar was driven at a frequency of 10 Hz.

#### 3) Experiment 3

This experiment is similar to experiment 1 with the difference that the joystick was removed and closed-loop brain control was employed to drive the wheelchair driven at a frequency of 10 Hz. A Linear Discriminant Analysis classifier based decoder was calibrated to achieve closed loop brain control, details of which can be found in [5].

#### 4) Experiment 4

In this experiment (see Fig. 2 (b)), NHP B was trained to perform the classical centre-out task [13], [26], [27] through joystick control with targets appearing in one of the 8 different locations. A cursor appeared at the centre of the screen at the beginning of every trial and a trial was considered successful if the NHP managed to manipulate the cursor to reach and stay in the target area for 2 seconds under a total elapsed time of 10 seconds. The cursor position was updated every 100 ms via joystick control.

### C. Pre-recorded dataset inputs and outputs

The number of spikes occurring in a backward looking window of time, *T_w_* = 500 ms, at each of the input *D* electrodes constitutes the input feature vector, 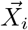, corresponding to the time, *t* = *i*,

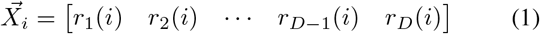

where 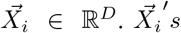 are determined in a sliding window fashion with a step-size of *T_s_* = 100 ms, for the entire duration of a trial across all the reported experiments. In case of supervised learning algorithms, a one-hot target vector 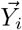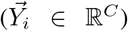 exists for corresponding 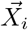, in order to enable learning. *C* stands for the number of discrete classes. In experiments 1, 2 and 3, *C* = 4 corresponding to joystick outputs - *left*, *right*, *forward* and *stop*. *C* = 8 corresponding to the eight different possible target locations (see Fig. 2(b)) in experiment 4. RL algorithms do not employ full output label information containing, 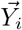. Instead a binary scalar feedback exists at every time-step, *t* = *i* to enable learning. Feedback signal at *t* = *i* is given as,

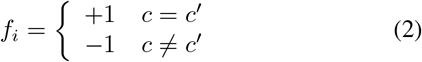

where *c* and *c′* are predicted and true labels respectively at the previous time-step, *t* = *i* − 1.

### D. Datasets and train-test split

We have analyzed four datasets acquired from two NHPs in an offline manner. In experiments 1 and 2, we have used a total of 8 days of recorded data with 3 sessions on each day. In experiments 3 and 4, we have used data recorded on 4 separate days with 3 sessions on each day.

Fig. 1(d) shows the manner of train-test data split. Test data is consistent across all algorithms and corresponds to the sessions 2 and 3 on recorded days, across all experiments. Training data corresponds to session 1 of each day for supervised daily *retrained* models, and session 1 of day 1 for supervised *fixed* models. RL algorithms do not employ explicit training data. They are randomly initialized at the beginning of a recorded day and are updated in a purely online manner on test data.

## III. Reinforcement Learning Algorithms

### A. Proposed

#### 1) Banditron

Initially at time *t* = 1, Banditron weight matrix is initialized as ***W***^**1**^ = **0**∈ℝ^*C×D*^, where *C* represents the number of output classes and *D* represents the number of input channels. At subsequent time-steps, *t* = *i*, as the iBMI system receives inputs, 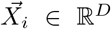, Banditron utilizes current weight matrix ***W** ^i^* to arrive at an intermediate output, 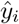,

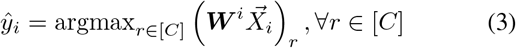

where [*C*] = 1*, …, c*.

Thereafter, for ∀*r* ∈ [*C*], Banditron defines a probability, *P* (*r*) over classes as,

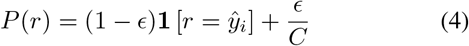

where **1**[*π*] is 1 if predicate *π* holds and 0 otherwise. The final predicted value, 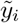, is randomly sampled according to probability *P* (*r*). This enables the algorithm to *exploit* current weight matrix (***W**^i^*) to choose 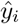 with probability 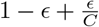, and *explore* with probability 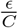 to randomly choose among all the *C* output labels. Thus, *ϵ* determines the exploration-exploitation tradeoff in Banditron. Suggested range for choosing *ϵ* is given as *ϵ* (0, 0.5).

Based on the feedback (1 if actual output, *y_i_* equals final predicted value, 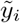 and 0 otherwise), the weight matrix is updated as,

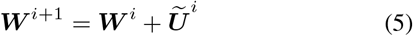

where 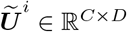 is the update matrix given as,

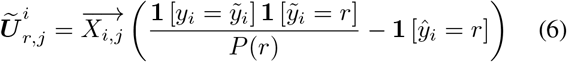

Do note that the true label (*y_i_*) is never revealed to the algorithm, but we indirectly obtain full information when, 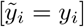.

#### 2) Banditron-RP

Banditron is inspired by the simple linear perceptron and hence it follows that it is limited in its capability to learn only linearly separable patterns [23]. However, superior performance has been reported by non-linear methods over linear ones in iBMIs [8], [28]. Thus, we have introduced non-linearity in the form of obtaining feature vector - 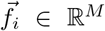 as a non-linear random projection (RP) of 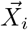 given as,

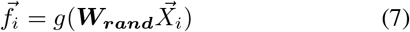

where ***W_rand_*** ∈ ℝ^*D×M*^ is the fixed random projection weights chosen from a standard uniform distribution in the interval (0,1), *g*(.) corresponds to the activation function which in our case is sigmoid. Learning proceeds in the same manner as Banditron with 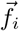 serving as input instead of 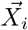. We henceforth refer to this variant of Banditron with input 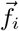 as *Banditron RP*.

Addition of a fully connected feature extraction layer to Banditron is bound to drastically increase the overall power dissipation. Thus, we propose implementing the first layer of fixed random weights, ***W_rand_***, in the form of single transistor based current multipliers reported in [10]. The custom designed chip - Spike-input Extreme Learning Machine (SELMA) [10] has the potential of directly being used to implement Banditron-RP.

### B. State of the art

#### 1) AGREL

Authors in [29] have introduced attention-gated reinforcement learning (AGREL) as a three-layer neural network which takes neural state as an input and maps it to corresponding action space. *j^th^* hidden layer node’s output is given as,

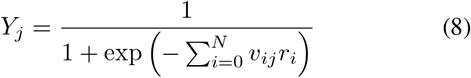

where *N* stands for input electrodes, *r_i_* is the spike firing rate appearing at the *i^th^* electrode, *v_ij_* represents the input layer weights and *r*_0_ = 1 to factor in input weight bias.

AGREL uses stochastic softmax rule to choose a winning neuron among *C* output neurons. Here *C* stands for the number of possible different actions. *k^th^* neuron’s output - *Z_k_* and the probability of choosing one among *C* actions is given as,

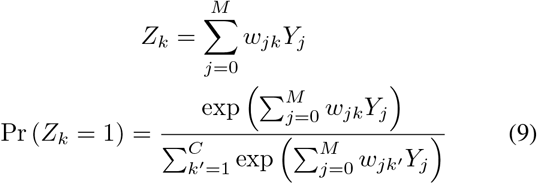

where *w_jk_* represents the output layer weights, *M* is the number of hidden nodes.

Instantaneous reward is defined as,

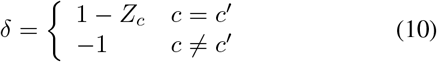

where *c* and *c′* are predicted and true actions respectively.

In order to expedite learning, AGREL defines an expansive function based on instantaneous reward as follows,

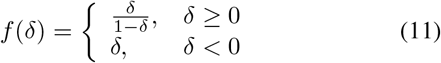

Weights are initialized randomly drawn from a standard uniform distribution in the interval (−1,1) [29]. At every iteration of prediction following the forward pass, weights are updated in the backward pass as follows,

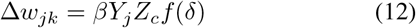

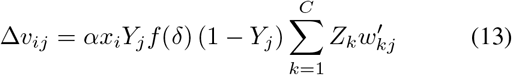

where *α* and *β* are learning rates.

#### 2) HRL

It uses a three-layer neural network akin to AGREL albeit with a difference in the way it goes about learning. Output of each hidden node (*OutH_i_*) is given as,

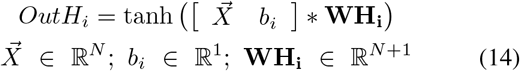

where 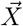 is the input feature vector, *b_i_* is a bias term and **WH_i_** stands for input weights.

Action value at the output nodes is given as,

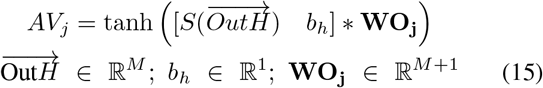

where *S*() stands for the sign function, *b_h_* is a bias term and **WO_j_** is the output weights. The output node with the highest action value among a total of *C* nodes is chosen to be the final output in this scheme. Feedback *f* is defined as,

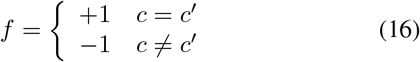

where *c* and *c′* are predicted and true actions respectively.

Initial weights are initialized randomly from a standard normal distribution [7] and are updated at every iteration with learning rates *μ_O_*, *μ_H_* in the following manner with action value output vector, 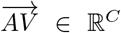,

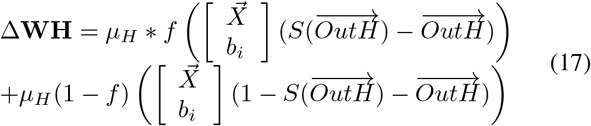

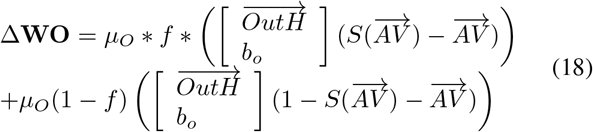

#### 3) Deep Q-learning

It involves a multi-layer neural network that takes the current state, *s* as an input variable and outputs Q values, *Q*(*s, c*) value for each possible action in the action space set [*C*] = 1*, …, c* [30], [31]. We have used a two hidden layer neural network implementation in the this study. Final output, *y*, is selected following the *E*-greedy policy as,

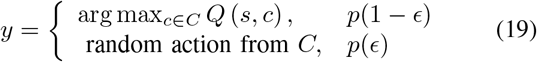

where *ϵ* determines the exploration-exploitation tradeoff. Q-learning involves the following update rule,

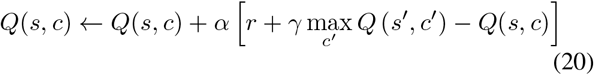

where *α* is the learning rate, *r* stands for the reward obtained when taking action *c* in state, *s*, *γ* is the discount factor and max_*c′*_ *Q* (*s′, c′)* is the maximum possible Q-value in the next state, *s′*.

Furthermore, the mean squared loss function to be minimized during the update phase is defined as,

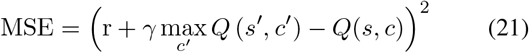

where r + *γ* max_*c′*_ *Q(s′, c′)* represents the target and *Q*(*s, c*) is the prediction.

## IV. Results

### A. Summary of models and Hyperparameter Tuning

Table I gives a summary of algorithms and their respective training paradigms analysed in this paper, whereas Table II captures the range of values and choices considered for arriving at optimal hyper-parameters for different algorithms. For SVM, we have used bayesian hyper-parameter optimization method *bayesopt* provided in Matlab R19a (The MathWorks, Inc., Natick, Massachusetts, United States) to tune hyper-parameters over 30 iterations in a five-fold cross-validation manner based on training set. LDA does not have any associated hyper-parameters. Hyper-parameters corresponding to the different RL algorithms are tuned on first recorded day’s session 1 across all datasets.

**TABLE I:** Summary of training paradigms and algorithms

|  | Supervised Learning | Reinforcement Learning |
| --- | --- | --- |
| Batch training, weight frozen | $LDA_{fixed}$,<br>$LDA_{retrain}$,<br>$SVM_{fixed}$,<br>$SVM_{retrain}$ | — |
| Only online, update | — | Banditron, AGREL,<br>HRL, Q-learning,<br>Banditron-RP |
| Batch training, online update (BTOU*) | — | $AGREL_{BTOU\_epochs\_xx}$<br>$AGREL_{BTOU\_transfer\_epochs\_xx}$ |

\*We explicitly refer to the RL methods with batch training followed by online updates with the subscript *BTOU*, whereas the RL algorithms with only online update have no subscript. *xx* in the subscript stands for number of training data replications (also known as epochs).

**TABLE II:**
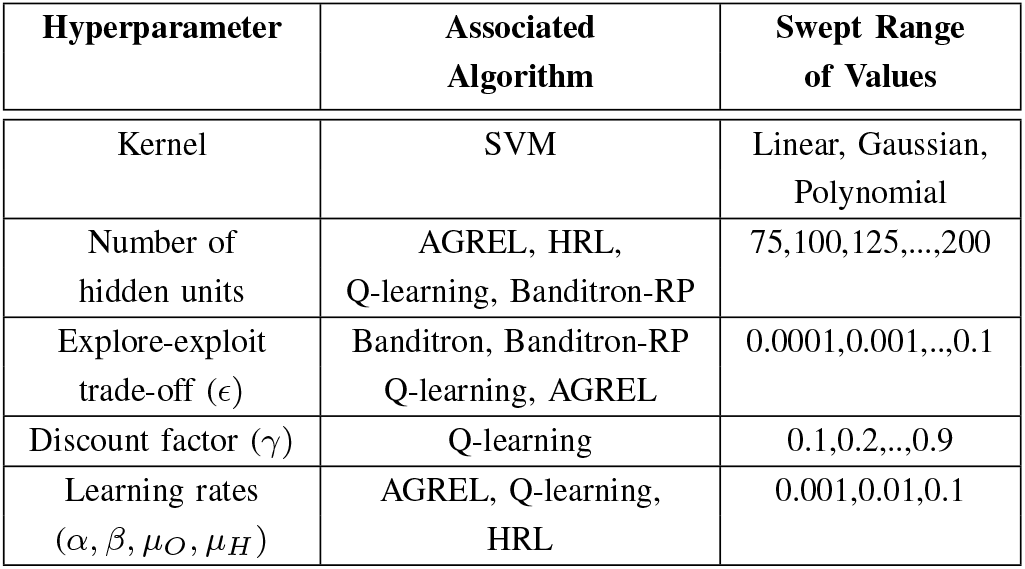
Hyperparameter tuning - parameter choices

### B. Decoding Results - Ideal feedback

We have reported classification accuracy as a measure of performance to compare decoding prowess of different techniques. Experiments 1, 2 and 3 represent a 4 class classification problem - *right*, *forward*, *stop* and *left*. Experiment 4 on the other hand has 8 distinguishable classes corresponding to the 8 different directions shown in Fig. 2(b).

Results in Fig. 3 show Banditron’s average improvements over AGREL, HRL and Q-learning of 6.01 ± 2.54%, 14.5 ± 1.83%, 4.83 ± 1.83% in experiment 1, 20.98 ± 14.95%, 14.97 ± 1.95%, 10.07 ± 11% in experiment 2, 6.7 ± 6.66%, 16.62±8.23%, 9.12±9.76% in experiment 3 and 7.44±2.72%, 25.04 ± 4.35%, 16.85 ± 2.53% in experiment 4 respectively. Statistical comparison of Banditron against AGREL, HRL and Q-learning yield p-values of 0.0078, 0.0078, 0.0078 in experiment 1 and 0.0391, 0.0078, 0.0391 in experiment 2 respectively for a Wilcoxon signed-rank test. This shows that the improvements afforded by Banditron are statistically significant (*p <* 0.05) over AGREL, HRL and Q-learning.

**Fig. 3:**
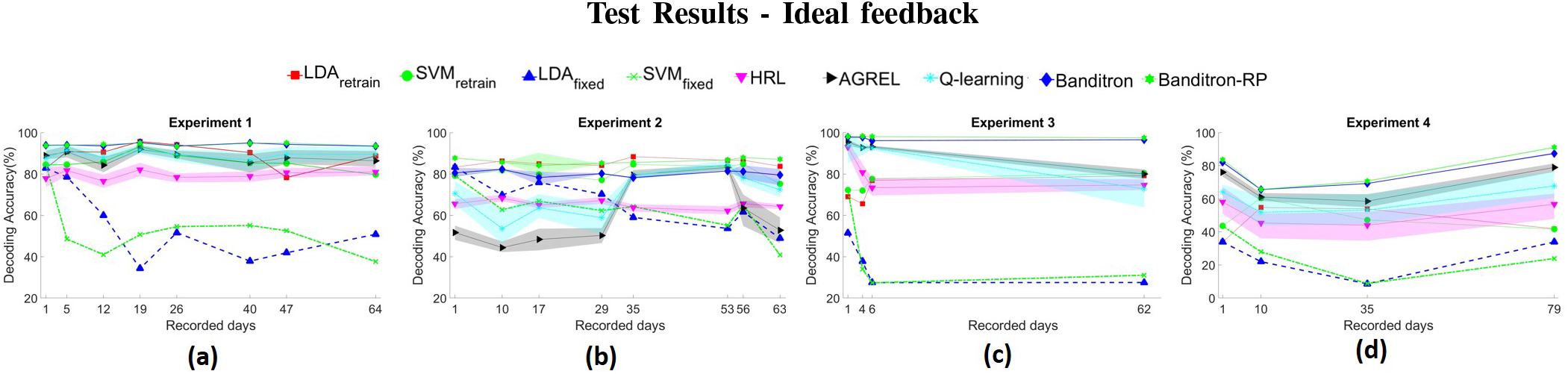
Decoding accuracy across four experiments has been reported for fixed and daily r etrained supervised models - LDA, SVM alongside RL algorithms - AGREL, HRL, Q-learning, Banditron and Banditron-RP. Shaded regions represent standard deviation of results across 20 iterations of random instantiations of probabilistic algorithms. Banditron and Banditron-RP significantly outperform the state of the art RL algorithms and *fixed* supervised models.

We have not reported p-values for experiments 3 and 4 as the number of data points are too less [32]. Furthermore, non-linear version of Banditron, Banditron-RP yields average improvement of ≈6% and 2% over Banditron in experiments 2 and 4 while performing as good as Banditron in experiments 1 and 3 respectively.

Meanwhile, Banditron yields substantial average improvements over fixed models *LDA_fixed_*, *SV M_fixed_* of 39.4 ± 18.39%, 41.06 ± 14.09% in experiment 1, 14.89 ± 11.66%, 18.19 ± 10.9%, in experiment 2, 61.02 ± 10.62%, 55.91 ± 20.31%, in experiment 3 and 51.51 ± 7.35%, 50.05 ± 13.86% experiment 4 respectively. Changes in recording conditions resulting from electrode impedance changes, micro-motion of electrodes etc. lead to non-stationary distribution of input neural data across days [4], [24]. This in turn leads to a drop in performance of fixed models - *LDA_fixed_*, *SV M_fixed_* due to the underlying shifts in data distribution [24], [28].

In case of comparison to the daily retrained supervised models, Banditron yields superior performance with average improvements over *LDA_retrain_*, *SV M_retrain_* of 5.21 ± 5.69, 8.19 ± 3.6 in experiment 1, 24.36 ± 7.17, 21.48 ± 4.89 in experiment 3 and 30 ± 19.65, 27.87 ± 18.16 in experiment 4 respectively. In experiment 2, Banditron performs almost as good as daily retrained classifiers with ≈94% level of performance.

### C. Decoding Results - Erroneous and sparse feedback

In Fig. 3, we have assumed the feedback at every time-step to be ideal corresponding to +1 for correct action and 1 otherwise along the lines of paradigm presented in [7], [33]–[35]. In order to study the impact of quality of feedback signals, we have added additional controls of, a) introducing error in feedback signals and b) making feedback sparse.

Errors are introduced in the feedback signal across 10% and 20% of the total time-steps on each day across all datasets. The erroneous time-steps on each day are chosen via a Matlab R19a (The MathWorks, Inc., Natick, Massachusetts, United States) function *randi* which chooses the fraction of erroneous time-steps from total time-steps following a uniform discrete distribution. Introducing error involves changing the feedback signal at a given time-step with value +1 to 1 and vice-versa. Furthermore, to study the impact of frequency of availability of feedback signal, we have reduced its occurrence to every second time-step (50% sparsity) and every fourth time-step (75% sparsity). Sparse feedback entails that the RL-decoder model is updated only when feedback is available. Figures 4 and *S*1 (in supplementary material) capture the results associated with erroneous and sparse feedback respectively.

**Fig. 4:**
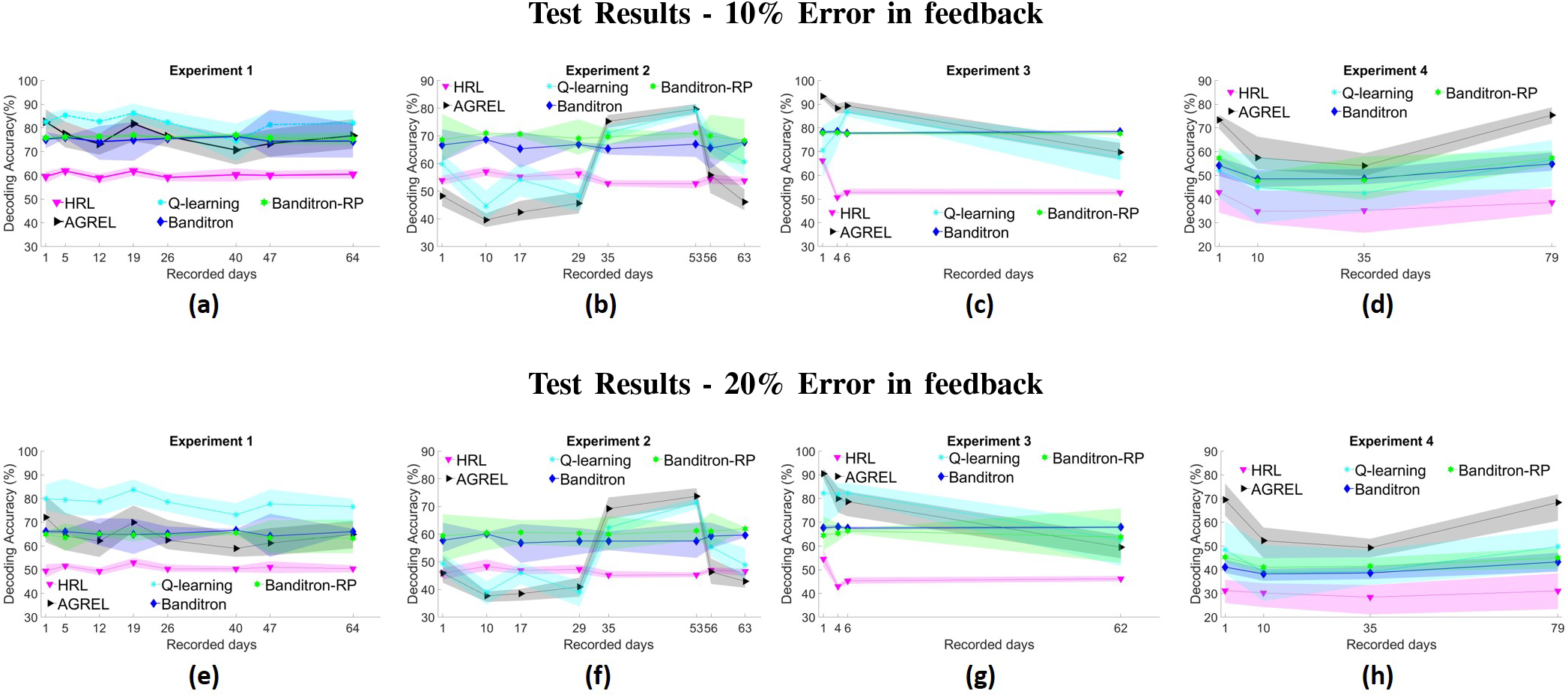
Decoding accuracy across four experiments has been reported for RL algorithms - AGREL, HRL, Banditron, Banditron-RP and Q-learning for 10% error in feedback in (a), (b), (c), (d) and 20% error in feedback in (e), (f), (g), (h) respectively. Banditron and Banditron-RP are susceptible to erroneous feedback and cease to outperform state of the art RL algorithms in all but experiment 2. Shaded regions represent standard deviation of results across 20 iterations of random instantiation of weights of probabilistic algorithms. Q-learning performs the best in experiment 1 while AGREL performs the best in experiments 3 and 4.

When deliberately introducing error in feedback signal, Banditron ceases to yield the best performance among RL algorithms in all but one experiment, experiment 2. In experiment 1, Q-learning performs the best with 6.96 ± 3.93%, 12.97 ± 3.48% average improvement over Banditron in 10%, 20% error cases. In experiments 3 and 4, AGREL performs the best with average improvements of 6.88 ± 10.67%, 9.4 ± 13.03% over Banditron in 10%, 20% error cases respectively in experiment 3 and average improvements of 13.52 ± 7.48%, 19.55 ± 8.55% in 10%, 20% error cases respectively in experiment 4. However, in experiment 2, Banditron still yields the best response over AGREL, HRL and Q-learning by 12.57 ± 15.69%, 12.12 ± 1.52%, 6.07 ± 12.13% in 10% error case and 8.87 ± 14.5%, 11.72 ± 1.1%, 6.84 ± 11.62% in 20% error case respectively. This goes to show that correct feedback is essential to Banditron yielding the best performance.

In case of introducing sparsity, Banditron still yields the overall best performance over other RL algorithms as seen in Fig. *S*1 in supplementary material. In case of 50%(75%) sparse feedback scenarios, Banditron achieves percentage improvements of at least ≈9%(11%), 11%(14%), 11%(13%), 6%(3%) in experiments 1, 2, 3 and 4 respectively over AGREL, HRL and Q-learning.

## V. Computational Complexity

We consider an iBMI system with *N* inputs for a *C* option discrete control. The number of multiply-and-accumulate operations (MACs) required during prediction for single layer classifiers such as LDA, Banditron can be given as,

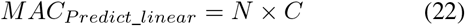

For single hidden layer neural network based approaches such as HRL, AGREL with *M* hidden nodes and a bias term in both input-hidden (*WH_i_*) and hidden-output weight (*WO*) matrices, the number of MACs required during prediction is given as,

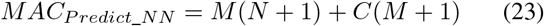

For Banditron-RP, we do not consider any bias term and accordingly, the number of MACs during prediction is given as,

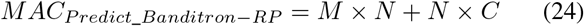

In Q-learning we have used two-hidden layers with *M*_1_ and *M*_2_ nodes respectively and corresponding bias terms. Accordingly, the number of MACs required during prediction step is given as,

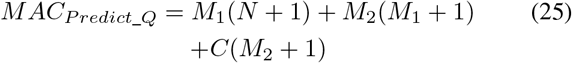

We have ignored the number of MACs required to implement the activation function for the sake of simplicity.

RL algorithms are recursively updated and MACs expended by single-layer Banditron and multi-layer neural network systems - AGREL, HRL, Q-learning and Banditron-RP can be formulated as below,

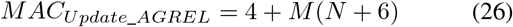

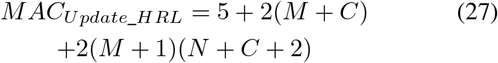

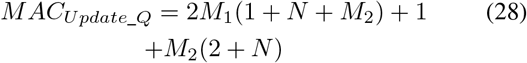

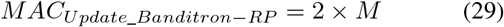

For a single layer Banditron based network, number of MACs required per update is given as

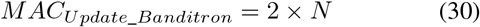

Do note that state of the art supervised schemes such as LDA, SVM are trained at the beginning of the day/session and are held fixed without being updated iteratively.

Consider a case of *N* = 64 input electrode iBMI system driving a 4-option (*C* = 4) discrete control at an operating frequency of 10 Hz, similar to the reported experiments in this paper. For neural networks based approaches HRL and AGREL, let us assume number of hidden nodes to be *M* = 80. To compute power dissipation we assume, Energy/(*Analog MAC*) = 0.45 *pJ* [10] and Energy/(*Digital MAC*) = 10 *pJ* [36]. With these assumptions, we can arrive at the Table. III comparing number of MACs memory requirement and estimate power dissipation across algorithms. Note that we have considered a linear version of SVM in Table. III. Furthermore, we consider weight resolution to be 16-bits across all algorithms.

**TABLE III:** Computational Complexity Comparison

| Algorithms | MACs update | MACs prediction | Memory (kB) | Power dissipation (nW) |
| --- | --- | --- | --- | --- |
| LDA, SVM | - | 256 | 0.5 | 25.6 |
| AGREL | 5604 | 5524 | 10.8 | 1112.8 |
| HRL | 11513 | 5524 | 10.8 | 1703.7 |
| Q-learning | 28481 | 12004 | 23.3 | 4048.5 |
| Banditron | 64 | 256 | 0.5 | 32 |
| Banditron-RP | 160 | 5440 | 10.6 | 560(71*) |

MAC = multiply and accumulate, Number of input channels, $N = 64$ , Number of hidden layer nodes, $M, M_1, M_2 = 80$ , Number of outputs, $C = 4$ , Weight resolution( $W_{res}$ ) = 16 bits. Memory = (MACs prediction) $\times W_{res}$ . Energy/(Analog MAC) = 0.45 pJ [10] and Energy/(Digital MAC) = 10 pJ at operating frequency, $f_{op} = 10$ Hz. \* reports estimated power dissipation for Banditron-RP implemented on custom chip - SELMA.

The values populated in Table. III show that Banditron is at least an order of magnitude less computationally complex in terms of number of MACs and memory required than neural network based RL algorithms - AGREL, HRL and Q-learning. This low complexity feature fits well with the prospect of implementing low power iBMI decoders [8], [37]– [39]. Furthermore, the number of MACs required by Banditron are in the same order of magnitude as LDA, SVM.

## VI. DISCUSSION

### A. Comparison to state of the art RL algorithms

#### 1) Effect of Variability in Neural Data

As observed in Fig. 3, the backpropagation (BP) based algorithms - AGREL, HRL, Q-learning perform poorly and erratically compared to Banditron, Banditron-RP. As an example of erratic performance, AGREL can be seen performing significantly better on days 35 and 53 in Fig. 3(b) compared to the other days. We hypothesize that intra-day nonstationarity/variability is responsible for this phenomenon, wherein variability makes generalization difficult for BP-based algorithms operating in the purely online mode.

In order to validate our hypothesis, we first compute the mean value of minimum principal angles (MPAs) across trials in a sequential manner on all days of experiment 2. MPA is a parsimonious scalar metric used to evaluate the level of similarity between neural datasets [28]. Lower values of MPA correspond to higher levels of similarity and vice-versa. Thus, the mean value of MPA across the sequence of trials serves as a metric of variability exhibited by the neural activity. In particular, we compute the MPA between the tuples - (*trial*_1_, *trial*_2_), followed by (*trial*_2_, *trial*_3_) until we reach the last trial tuple - (*trial_last−_*_1_,*trial_last_*)). Thus, our variability metric, *V ar.*, for total trials, *N_trials_*, can be given as,

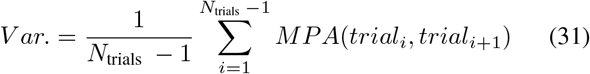

Days 35 and 53 as seen in Fig. 5 exhibit relatively lesser intra-day variability, thereby leading to improved performance for AGREL.

**Fig. 5:**
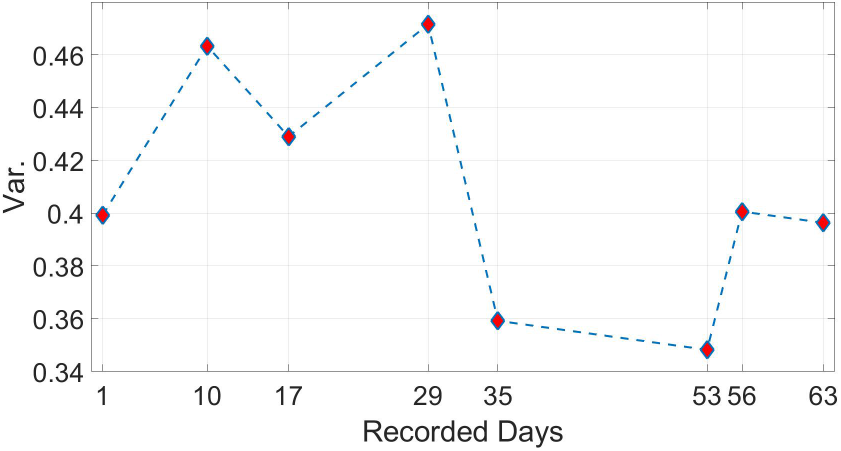
Shows intra-day variability of firing rates of trials across days in experiment 2. Variability is captured in terms of the mean value of minimum principal angles (MPAs) computed between trials on each day. Days 35 and 53 have relatively low variability compared to the other days.

In order to further validate the hypothesis about variability making generalization difficult for BP-based RL algorithms, we simulated two synthetic neural datasets, *Syn Dir* 4 and *Syn Dir* 8 along the lines of [40] using the Izhikevich model [41] for neural excitability. *Syn Dir* 4 and *Syn Dir* 8 correspond to the synthetic neural spike data generated for four-options (experiment 2) and eight-options (experiment 4) cursor control respectively. The detailed simulation methodology is reported in Section I of supplementary material. We compared the decoding performance of RL algorithms against varying percentage of noisy neurons in Fig. 6 for these datasets. All the algorithms were operated in the only online update mode. The inset figure in Fig. 6 represents percentage of noise added along x-axis and variability referred to as Var. (mean value of minimum principal angles across trials sequence – Equation 31) along the y-axis.

**Fig. 6:**
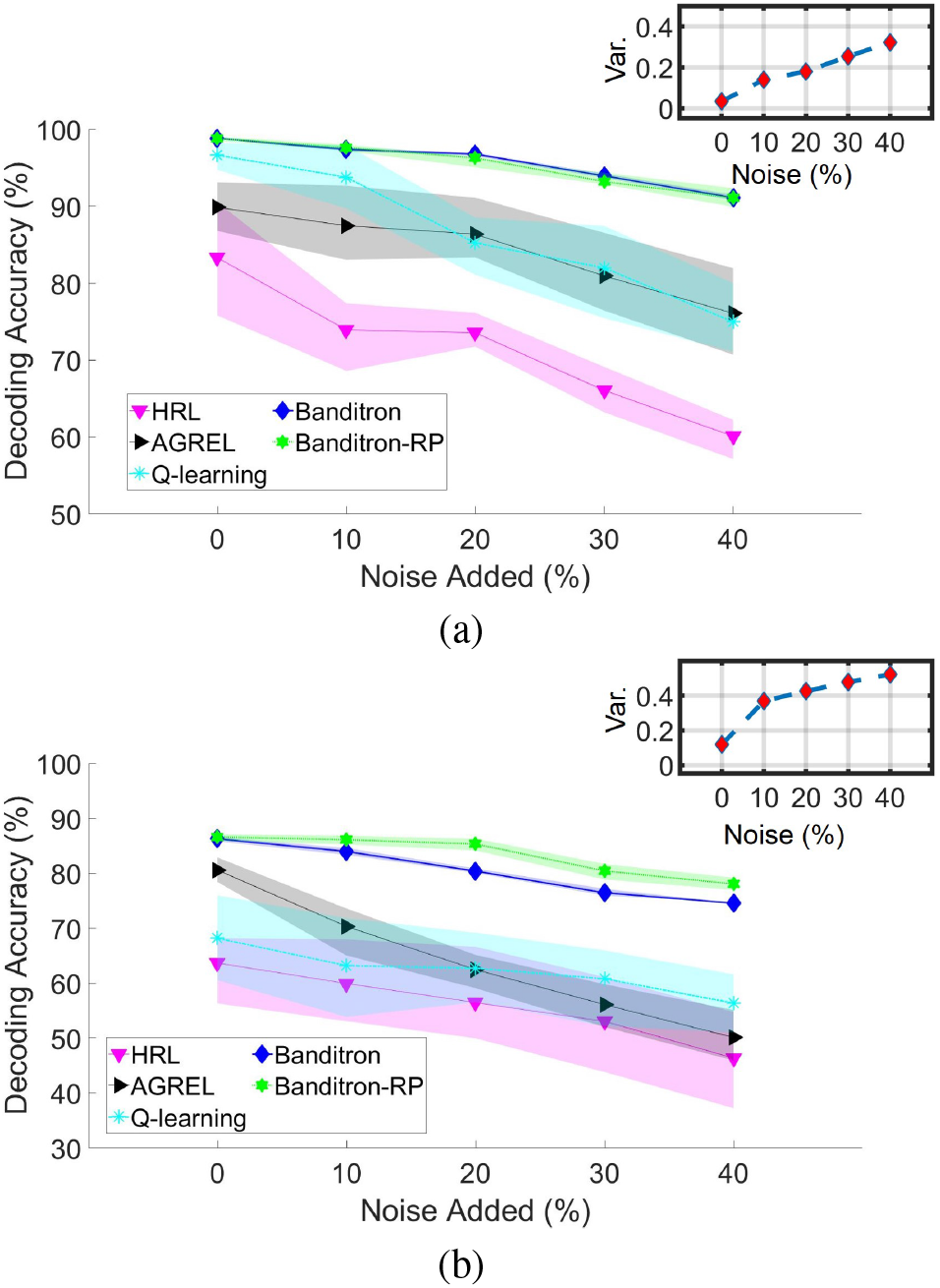
Shows decoder results for synthetic datasets (a) *Syn Dir* 4 and (b) *Syn Dir* 8 respectively across RL algorithms operated in the only online update mode. The figure in inset represents the Var. (Equation 31) on y-axis plotted against percentage added noise on x-axis. Variability increases as we add noise. Decoder accuracy is seen to be decreasing with increase in added noise, and at faster rate for BP-based RL algorithms.

The general trend observed in Fig. 6 is that the decoding performance of RL algorithms decreases with increase in variability. However, one must note that BP based RL algorithms such as HRL, AGREL and Q-learning suffer a relatively larger decline compared to Banditron, Banditron-RP. These results are qualitatively similar to the earlier presented results in Figures 3(b) and 5 wherein BP-based RL algorithms generalized relatively worse on days with more variability and vice-versa. Thus, the results in Fig. 6 validate our previously stated hypothesis in a controlled simulation setting. We observe that non-stationarity in neural data makes it harder for BP-based RL algorithms to generalize in an online update mode. However, simple linear Banditron and its nonlinear projection variant Banditron-RP perform better due to its fewer trainable parameters.

#### 2) Batch Training Mode

NNs are known to for their ability to approximate arbitrary functions following the universal approximation theorem. However, one must note that NN-based approaches trained by BP typically generalize well when trained over multiple replications of training data (also known as training epochs) [17], [42]. Keeping this in mind, we consider an alternative more conventional batch-based setting involving a dedicated set of training data for a BP based algorithm. To carry out this analysis, we consider AGREL as the BP based algorithm and use experiment 2 dataset. We name the NN model as *AGREL_BT OU epochs xx_*, where *BTOU* stands for batch training, online update, and *xx* stands for the number of training epochs. We train *AGREL_BT OU epochs xx_* on experiment 2’s - session 1 of each day (similar to daily retrained supervised models) over 5, 10 and 20 training epochs (replications of training data).

Thereafter, we deploy it as a decoder on sessions 2 and 3 on each day. We let it update sequentially in an online fashion as the binary feedback is received. In this case, we observe *AGREL_BT OU epochs xx_* comfortably outperforming the online version of AGREL. Thus, having a dedicated training session comes with the prospect of improved decoding performance. However, this comes with the cost of significant amount of calibration time [11]. As mentioned earlier, the quantum of calibration time amounts to approximately one-third of the total experiment time in our reported experiments.

In Fig. 7, we can see a saturation in performance of *AGREL_BT OU epochs xx_* after 10 epochs. Similar convergence can also be seen in supplementary Fig. *S*2, where we show a plot of training (session 1) and validation accuracy (sessions 2 and 3) across number of epochs for day 1 of experiment 2. This shows that no further improvement is expected by increasing the number of epochs and we have trained the network optimally.

**Fig. 7:**
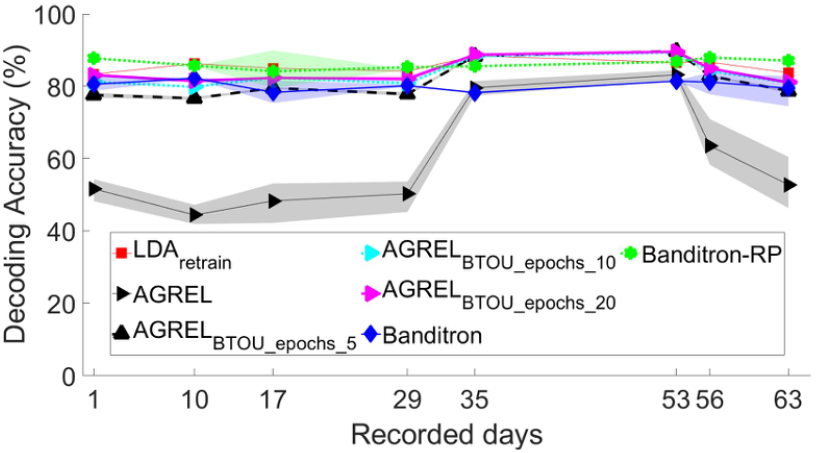
A more conventional batch-based train-test setup showing performance of an BP based RL algorithm, AGREL, when trained on a dedicated training set over 5, 10 and 20 training data replications (epochs) on experiment 2 dataset. The version employing a dedicated training set is referred to as *AGREL_BT OU epochs xx_*, where *xx* stands for the number of epochs. *AGREL_BT OU epochs xx_* comfortably outperforms online AGREL while closely matching *LDA_retrain_*’s results as we increase the number of epochs to 10 or higher.

As observed in Fig. 7, having a dedicated training session, leads to *AGREL_BT OU epochs xx_*’s performance to reach approximately the level of supervised learning algorithm – *LDA_retrain_*, as we increase the number of epochs. This is in consonance with the universal approximation view, and we expect to achieve qualitatively similar results for other NN based approaches in other experiments as well. However, do note that the 2 layer version of Banditron with randomized neurons in the first layer (referred to as Banditron-RP) also achieves similar performance as *AGREL_BT OU epochs_* 10 while retaining the fully online version of training. Hence, this seems like a good choice to balance network learning capacity with online learning for iBMI applications.

A point to note here is that *AGREL_BT OU epochs_* 10 also outperforms online version of Banditron. This shows that in the event of a dedicated calibration procedure NN-based models such as AGREL outperform simple online linear model - Banditron. However, this superior performance comes at the cost of calibration times and computational complexity. If one were to implement training on chip, Banditron provides savings of at least 6 orders of magnitude in MACs and memory requirement compared to NN models. In addition to the assumptions listed in Table. III, we have assumed calibration over 10 epochs over recording time of 5 minutes with a time-step *T_s_* = 100 *ms* and input firing rate resolution *FR_res_* = 8 *bits*.

#### 3) Transfer Learning

A common solution to paucity of data is to train a NN on a related dataset and thereafter update only the last few layers following the paradigm of transfer learning popularly employed in both computer vision [43] and in biomedical applications [44]. The hope is that the model learns complex feature representations that can be easily deployed on similar tasks.

Following this approach, we performed a preliminary experiment on experiment 2 dataset with a single hidden layer *AGREL_BT OU transfer epochs_* 10 model trained on session 1 of each day over 10 epochs. The number of training data epochs was set based on the results observed in Fig. *S*2 in supplementary material, wherein saturation in performance was observed at 10 epochs. Thereafter, we fixed the first layer weights and updated only the last layer weights for incoming samples from test sessions 2 and 3. We used the Banditron single layer rule to update the output layer weights. We observed however (Fig. 8) that the learned features do not yield any improvement in decoding performance over random features used in Banditron-RP again pointing to variability in input data statistics.

**Fig. 8:**
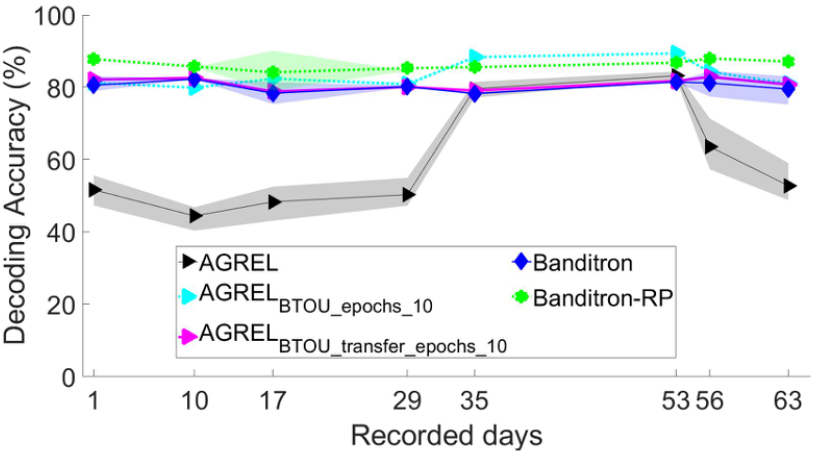
Applying transfer learning, while using an AGREL-trained NN as a feature extractor on experiment 2 dataset. We train the model *AGREL_BT OU transfer epochs_* 10 on 10 epochs using session 1 as training data. Subsequently, we fix the first layer weights and update the last layer weights using Banditron weight update rule. *AGREL_BT OU transfer epochs_* 10 does not yield improvements over simple Banditron.

In order to investigate the lack of improvement in performance, we visualize a plot of low dimensional projection of input firing rates and extracted features for training (session 1) and test data (sessions 2 and 3) across two days for experiment 2 in Fig. *S*3 (supplementary material). The extracted feature embedding corresponds to the hidden layer of the aforementioned model - *AGREL_BT OU transfer epochs_* 10. Please note that for the purpose of ease of visualization, we showed only three clusters instead of the original four employed in the experiment. Visual inspection shows that the separability looks better for the extracted features on training set as seen in Fig. *S*3 (a),(b) in supplementary material. However, separability looks somewhat similar on test data as seen in Fig. *S*3 (c),(d) in supplementary material. This, explains why the performance of *AGREL_BT OU transfer epochs_* 10 acting on extracted features is roughly equivalent to Banditron acting on input features.

### B. Comparison to state of the art supervised learning paradigm

Compared to supervised retrained models, Banditron reports similar (experiment 2) or better (experiments 1, 3 and 4) levels of performance as *LDA_retrain_*, *SV M_retrain_* (see Fig. 3). Supervised learning corresponds to the state of the art decoding methods employed in iBMIs. Thus, we have employed popular techniques - Linear Discriminant Analysis and Support Vector Machines to establish a baseline for the RL based results. One must note two stark differences between supervised learning and RL algorithms (see Fig. 1(a)) - a) Supervised algorithms employ an explicit training phase, whereas RL algorithms do not have one and b) Supervised algorithm based models employed in iBMIs typically remain fixed once trained, whereas RL algorithms are constantly updated as they interact with the environment. Thus, this comparison should be seen as a way to establish context in relation to state of the art techniques and not as a one-to-one comparison.

### C. Towards Autonomous iBMIs

The online continuous adaptive nature of the reported paradigm of iBMI operation coupled with possible biological sources of reward signal lead us to envision an autonomous iBMI system wherein no explicit calibration trials are required and the system simply learns from the feedback signal. Thus, this paradigm has the potential to drastically reduce the involvement of the neuro-engineer in enabling day to day usage of iBMI by patients.

However, this requires the presence of a valid biological reward/error signal. In this study, we are using binary evaluative feedback signal at every time-step along the lines of works [7], [33]. The reason we have opted for this approach is that recent works have shown this kind of signal to be present in regions of the brain such as nucleus accumbens [20], primary motor cortex [21], [34], [35], [45]–[48]. Apart from these regions, other candidate sources of obtaining this signal include EEG [49], ECoG [50] among others. We certainly feel that more work is needed in this area to fetch a reliable and stable feedback signal.

An alternative to sourcing the reward signal from a biological source is to derive it from the effector’s trajectory as the agent attempts to move it towards a target [19]. In this approach [19], action chosen by the agent is rewarded if it moves towards the target and punished otherwise. This approach, while not useful for day to day autonomous usage by a patient, will still be useful in conducting iBMI experiments where the ground truth target is known from experimental design. We have reported open-loop results and it will be interesting to compare supervised approaches such as ReFIT [13] against RL-based approaches [19] in a closed-loop brain control setting.

## VIII. Conclusion, Limitations and Future Work

We have presented decoding results across different RL algorithms in a binary evaluative feedback scenario. Bandit algorithms – Banditron and Banditron-RP, for training a single layer of weights, are seen to be the best performing candidates among compared algorithms in an ideal feedback setting. Furthermore, they are seen to be relatively more robust to sparse feedback (see Fig. *S*1 in supplementary material), i.e. when a feedback signal is not available at all time steps as is likely going to be the case in real BMI scenarios. However, one must point out that Banditron based learning system’s superior performance is subject to accuracy of feedback. As seen in Fig. 4, it is observed that Banditron’s performance is not the best compared to other algorithms when feedback signals are erroneous. This implies a better strategy would be to estimate confidence in feedback signal first before updating weights using Banditron–this is an aspect for future in-depth study.

As far as the power dissipation is concerned, Banditron has the potential to offer at least two orders of magnitude in savings compared to state of the art RL algorithms. This has the potential to achieve a fully implantable solution and prolong the battery life of the iBMI.

Results presented in *Section* IV show that the nature of adaptive learning of this paradigm is naturally robust against non-stationarities. The current open-loop offline results hope to serve as a stepping stone towards closed-loop studies employing RL algorithms. Through the reported results we urge the community to advance along RL lines of research instead of simply focussing on supervised learning paradigm. Following prior works, we have employed a feedback signal at every time-step. However an interesting question we wish to ask in future studies is that, “Can the iBMI system still perform well if the system gets delayed reward only at the end of a trial (attempted movement)?”.

## ACKNOWLEDGMENT

The authors would like to thank Abdur Rauf and Clement Lim for helping in training the NHPs and data collection. This work was supported through grant RG 87/16 by MOE, Singapore.

