## supplementary material for "Lightweight Reinforcement Algorithms for autonomous, scalable intra-cortical Brain Machine Interfaces"

Shoeb Shaikh, *Student Member, IEEE,* Rosa So, *Member, IEEE,* Tafadzwa Sibindi, Camilo Libedinsky and Arindam Basu, *Senior Member, IEEE*

### I. Neural Dataset Simulation Methodology

$Syn\_Dir\_4$ corresponds to the synthetic neural spike data generated for four-options cursor control reported in experiment 2. We took experiment 2 - day 1's target matrix and obtained a neural matrix, $\boldsymbol{X}$ , ( $\boldsymbol{X} \in \mathbb{R}^{T \times N}$ , $T$ - number of time-steps and $N$ – number of input neurons) following the Izhikevich method given by,

$$v' = 0.04v^2 + 5v + 140 - u + I \tag{1}$$

$$u' = a(av - u) \tag{2}$$

with resetting of auxiliary spike,

$$\text{if } v \geq +30\text{mV then } \begin{cases} v \leftarrow c \\ u \leftarrow u + d \end{cases} \tag{3}$$

$v$ is the membrane potential and $u$ is the membrane recovery variable. $a = 0.02$, $b = 0.2$, $(c, d) = (65, 8)(15, 6) \cdot e^2$ where $e$ is a random variable uniformly distributed, $e \in [0, 1]$. $I$ is

$I$ takes on the value of 1 for a neuron tuned to a specific output action (target) or 0 otherwise at every time-step, $t = i$, for the tuned ensemble of neurons. The target value corresponding to every time-step is taken from day 1 of experiment 2. One must note that real world neural data suffers from issues such as electrode deterioration, electrode micro-motion, changes in electrode impedance among others. To account for these effects, authors in [1] propose changing value of a tuned neuron's 1 from $1/0$ to a value chosen at random from standard Gaussian distribution at every time-step, $t = i$. The proportion of such noisy neurons were added in steps of 10% from 0 to 40% to the tuned neuron ensembles in order to create five copies of neural spike data with varying degrees of noise [1]. the synaptic current which is calculated from target variable as 1 for spike and 0 for all other times.

Neural spike dataset - $Syn\_Dir\_4$ was generated for four target states corresponding to the four actions – left, right, forward and stop. We considered the number of input neurons to be $N = 60$, and split them into five ensembles comprising of 12 neurons each. Following the methodology reported in [1], we tuned four ensembles to each of the four output actions, and the fifth ensemble was left uncorrelated with the output action space. The fifth ensemble served to simulate noise and the synaptic current $I$ randomly chose values from a standard Gaussian distribution for these neurons.
Added noise introduces variability/non-stationarity in neural data.

Similarly, we created $Syn\_Dir\_8$ for eight output actions corresponding to movement towards the eight center-out targets. In this case, we used target matrix corresponding to day 1 of experiment 4 to arrive at the neural data matrix. We used $N = 63$ input neurons in this case and split them into nine equal sized ensembles – eight ensembles tuned to each of the eight directions and the remaining one for noise. Furthermore, we introduced additional noise in 0 to 40% of tuned neurons in steps of 10%, by changing the value of synaptic current from $1/0$ to a random value chosen from standard Gaussian distribution. This yields us five versions of neural spike data with varying degrees of noise (variability/non-stationarity).

**Test Results - 50% Sparse Feedback**

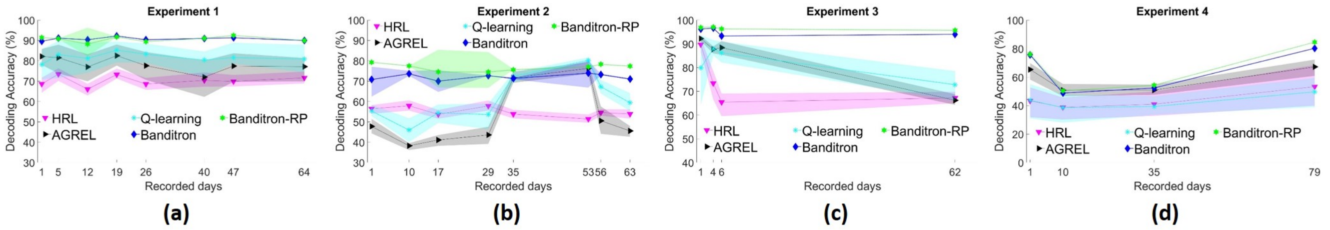


**Test Results - 75% Sparse Feedback**

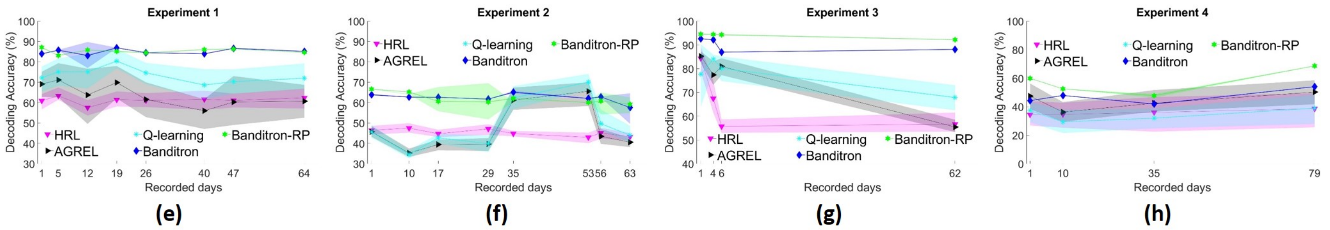


Fig. S1: Decoding accuracy across four experiments has been reported for RL algorithms - AGREL, HRL, Banditron, Banditron-RP and Q-learning withholding feedback signal across time-steps thereby introducing sparsity in feedback. (a), (b), (c), (d) depict results for $50\%$ sparsity in feedback and (e), (f), (g), (h) depict results for $75\%$ sparsity in feedback respectively. Shaded regions represent standard deviation of results across 20 iterations of random instantiations of probabilistic algorithms. In this scenario, Banditron and Banditron-RP significantly outperform the state of the art RL algorithms.

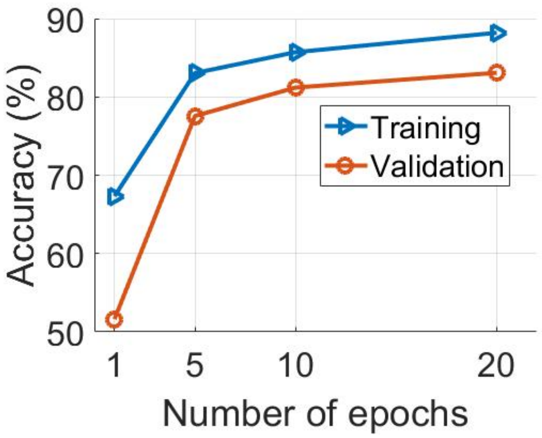


Fig. S2: This plot shows $AGREL_{BTOU\_epochs\_xx}$'s training accuracy and validation accuracy on experiment 2 on day 1 for varying number of training data replications (epochs). The improvement in performance saturates roughly after 10 epochs.

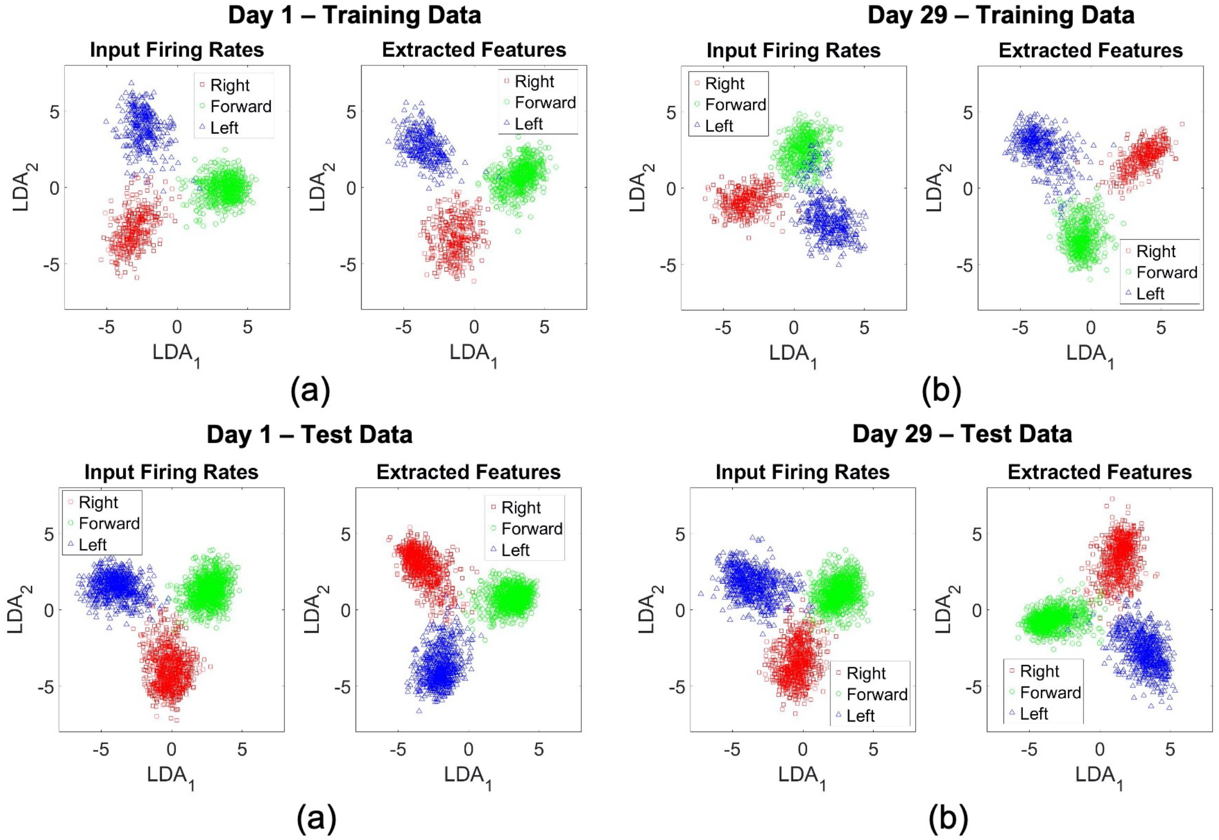


Fig. S3: Low dimensional representation of input firing rates and extracted features for - (a) day 1 and (b) day 4 of training data (session 1), and (c) day 1 and (d) day 4 of test data (sessions 2 and 3) corresponding to experiment 2 dataset respectively. $AGREL_{BTOU\_transfer\_epochs\_10}$ is used to learn complex feature representations following the paradigm of transfer learning. These representations are referred to as extracted features. The separability of extracted features appears relatively better than input firing rates in the training set, whereas no improvements can be observed in the testing set. Please note that we have only shown clusters corresponding to three options instead of the original four in the experiment for ease of visualization.
